# Deep learning-based tumor microenvironment segmentation is predictive of tumor mutations and patient survival in non-small-cell lung cancer

**DOI:** 10.1101/2021.10.09.462574

**Authors:** Łukasz Rączkowski, Iwona Paśnik, Michał Kukiełka, Marcin Nicoś, Magdalena A. Budzinska, Tomasz Kucharczyk, Justyna Szumiło, Paweł Krawczyk, Nicola Crosetto, Ewa Szczurek

## Abstract

Despite the fact that tumor microenvironment (TME) and gene mutations are the main determinants of progression of the deadliest cancer in the world – lung cancer – their interrelations are not well understood. Digital pathology data provide a unique insight into the spatial composition of the TME. Various spatial metrics and machine learning approaches were proposed for prediction of either patient survival or gene mutations from these data. Still, these approaches are limited in the scope of analyzed features and in their explainability and as such fail to transfer to clinical practice. Here, we generated 23,199 image patches from 55 hematoxylin-and-eosin (H&E)-stained lung cancer tissue sections and annotated them into 9 different tissue classes. Using this dataset, we trained a deep neural network ARA-CNN, achieving per-class AUC ranging from 0.72 to 0.99. We applied the trained network to segment 467 lung cancer H&E images downloaded from The Cancer Genome Atlas (TCGA) database. We used the segmented images to compute human interpretable features reflecting the heterogeneous composition of the TME, and successfully utilized them to predict patient survival (c-index 0.723) and cancer gene mutations (largest AUC 73.5% for *PDGFRB*). Our approach can be generalized to different cancer types to inform precision medicine strategies.

## Introduction

In clinical practice it is common to diagnose cancer based on hematoxylin-eosin (H&E) stained images obtained through biopsy or surgery^1^. Such images are routinely stored for each patient, so there is an abundance of patient-specific, disease progression-relevant data that until recently has not been utilised at scale in cancer research. This has changed with the advent of digital pathology and the development of machine learning approaches to various predictive tasks based on H&E image data^2,3^.

In addition to tumor cells, H&E images portray the spatial architecture of the tumor microenvironment (TME), including stromal cells, immune cells, and hypoxic/necrotic tissue areas and their reciprocal spatial arrangement. The TME plays an important role in cancer progression and metastasis, and thus it is critical to study its composition extensively^4^. Different tumors, even of the same type, have various genetic profiles resulting from gene mutations^5^. For a given cancer type, survival of individual patients can largely vary^6,7^. Finally, the TME of different tumors is also different^8^. A burning question in this context is how the structure of the TME relates to patient survival and gene mutations. This question is particularly relevant for lung cancer. Lung cancer is the deadliest cancer type worldwide, and lung adenocarcinoma (LUAD) is becoming the mainly diagnosed subtype of lung cancer^9^. LUAD includes a relatively higher proportion of cases without tobacco exposure. Thus, it has a more balanced molecular background and is more frequently associated with the presence of single somatic driver mutations that may be effectively managed with specific molecularly targeted therapies^10^. Several genes are known markers of response to treatment and survival in LUAD, including *EGFR, ALK, ROS1, BRAF, NTRK1-3, RET, MET, KRAS*, and diagnostic panels for targeted gene sequencing for detecting mutations in critical genes are routinely used in clinical practice^11^. H&E images are commonly inspected for LUAD diagnosis and prognosis^12^. Wide tumor spread, access to vessels, large areas of necrosis visible in H&E images, are associated with poor diagnosis^4^, while abundance of immune cells indicates anti-tumor response of the immune system and associates with better survival^13,14^. The TME plays an important role in LUAD response to immunotherapy. Expression of PD-1 and PD-L1 on cancer or immune cells, as well as tumor mutation burden (TMB), are important biomarkers of immune checkpoint inhibitor efficiency^15,16^. The interconnections between the spatial TME composition, gene mutations and LUAD patient survival are so far not well understood.

Computational prediction of patient survival from H&E images has been either performed based on spatial metrics or using deep learning approaches. In the former case, spatial metrics are initially used to summarize the spatial arrangement of different tissues and next their correlation with survival is investigated. Alternatively, these spatial metrics are used as features in traditional machine learning algorithms^17–20^. The TME can be very heterogeneous, so it is not obvious how to quantify it and what metrics to use. These metrics include proportion-based^21^, clustering-based^22^, and methods borrowed from ecology^23^, and they have been applied to many different cancer types^18,22,24–26^. In general, all these metrics share a common trait, i.e., they incorporate only a limited number of tissue types at once, such as tumor cells and lymphocytes, tumor cells and stroma, etc. This approach cannot comprehensively capture the complexity of the TME. Thus, there is an unmet need for an encompassing spatial metric that would consider many possible TME components at once. In the latter case, deep neural networks are trained to predict patient survival directly from H&E images. Such deep learning-based methods are increasingly used for survival prediction and have been shown to perform comparably to or even better than spatial metric-based approaches^27,28^. However, one major disadvantage of deep learning methods is the lack of explainability. Due to the complicated structure of these models and number of parameters, it is not easy to surmise which parts of the TME are the most important for patient survival. This creates a need for explainable H&E image-based survival models.

Numerous methods for predicting gene mutations from H&E images were introduced and applied to a spectrum of cancers^29–34^, showing that such approaches can reveal links between the TME composition and mutations of selected genes. However, similarly to deep learning-based methods for patient survival, these models take raw image data as input and directly predict the presence of mutations. As such, it is hard to assess what parts of the TME are most predictive of a given mutation. A recent study proved that human-interpretable features extracted from images segmented with deep learning methods can be successfully applied to predict phenotypic expression^35^. This suggests that the same approach can be implemented to predict patient survival and gene mutations, which has not yet been explored. To address these shortcomings, here we develop a framework for predicting survival and gene mutations in LUAD patients based on H&E images and using human-interpretable features. First, we train a deep learning classifier and apply it to segment the H&E images from 467 tumor LUAD samples into nine tissue classes. Next, we compute two human-interpretable spatial features that describe the composition of the TME in segmented H&E slides. Finally, we use these features in combination with clinical data to predict patient survival, as well as to predict tumor mutations. The predictions generated by our model are human interpretable, i.e., it is possible to pinpoint exactly which component of the TME is associated with a change in survival hazard or with a given mutation. Our framework is generalizable and can be readily extended to other tumor types to make predictions of patient survival and cancer mutations based on digital pathology. This work is a step forward to a better understanding of the interplay between the TME, gene mutations and survival of LUAD patients.

## Materials and methods

### Clinical samples

We obtained the formalin-fixed paraffin embedded (FFPE) tissue samples from 55 primary tumors of lung cancer (35 lung adenocarcinoma, 20 lung squamous cell carcinoma). The material was derived from FFPE surgical materials as well as from diagnostic (biopsy) procedures performed at Medical University of Lublin, Poland. In the moment of diagnosis and surgical resection of the primary cancer lesions, all patients were chemo-, radio-, immune- and targeted therapy naïve. We collected clinical and demographic patients’ data in a manner that protected their personal information. The study protocol received ethical approval from the Ethics Committee of the Medical University of Lublin, Poland (no KE-0254/235/2016).

### Extraction and annotation of the training dataset for ARA-CNN

We extracted the training dataset from the H&E slides sourced from 55 lung cancer patients. We loaded these slides into the QuPath software package^36^ and regions of contiguous tissue were annotated by an expert pathologist with the following nine labels: tumor, stroma, mixed, immune, vessel, bronchi, necrosis, lung, background. The annotated regions were then traced by a moving window, which cut out non-overlapping square patches of tissue with side size of 87 μm. In addition to 87 μm, we also tested the training performance for patches with sizes of 74 μm and 100 μm (see **Sup. Table 1-3**). They resulted in worse performance (mean accuracy 84.64% for 74 μm, 84.35% for 100 μm, 85.21% for 87 μm), so we proceeded with using the 87um sized ones. This gave us an initial version of the training dataset, which was then improved upon by utilizing human-in-the-loop active learning, as part of the previously proposed accurate, reliable and active (ARA) image classification framework^37^. In total, we ended up with 23199 patches, divided in the following manner: 3311 tumor patches, as well as 1511 stroma, 716 mixed, 1196 immune, 1236 vessel, 2030 bronchi, 4448 necrosis, 6031 lung, and 2211 background patches.

### ARA-CNN model

The main component of the ARA framework is ARA-CNN, a Convolutional Neural Network (CNN) architecture inspired in part by Microsoft ResNet^38^ and DarkNet 19^39^. It includes standard techniques for such models: Batch Normalisation^40^ (for normalisation and to reduce overfitting) and dropout^41^ (reduces overfitting). The latter allowed us to apply variational dropout^42^ during testing. Variational dropout is used to estimate uncertainty for every input image, which is returned together with its predicted class.

The model was trained in several iterations, each one improving upon the previous ones. After the first training process, the distribution of uncertainty for images in each class was measured separately and due to higher median uncertainty, it was concluded that there are three classes in need of more training examples: mixed, vessel and bronchi. These were passed on to the pathologist, who labeled new regions belonging to these classes. The resulting new training samples were extracted and added to the previous training dataset. This adaptive training procedure was repeated three times, where each time the uncertainty for each class was measured, until it was decided that the uncertainty results were at a satisfactory level.

For training the final ARA-CNN model on the lung cancer tissue patches, we used stratified 10-fold cross-validation. In each fold, the whole dataset of 23199 images was split into a training dataset and a test dataset used for evaluation. The test dataset contained 2316 patches, while the training dataset consisted of 20883 patches. Each class was split in exactly the same proportion: 10% were sent to the test dataset and 90% to the training dataset.

Additionally, in each training epoch the training data was split into two datasets: the actual training data and a validation dataset. The latter was used for informing the learning rate reducer - we monitored the accuracy on the validation set and if it stopped improving, the learning rate was reduced by a factor of 0.1. This split was in proportion 90% to 10% between actual training data and the validation set, respectively.

For parameter optimisation, we used the Adam^43^ optimiser. The training time was set to 100 epochs. The training data was passed to the network in batches of 32, while the validation and test data was split into batches of 128 images. The loss function used during training was the categorical cross-entropy.

### TCGA image data segmentation using ARA-CNN

The normalized patches (**Supplementary Methods**) served as input to ARA-CNN. For each input patch, the model returned a classification probability into each of the nine predefined classes. With these results, each patch was labeled with the class with the highest probability and then the labeled patches were merged back into their full respective slides and colored by the label. This created segmented slides, with clearly visible continuous areas of differing tissue.

The segmented slides were also validated by an expert pathologist, who assessed that 39 slides needed to be excluded from further analysis. There were two reasons for that. The first one involved erroneous classifications returned by ARA-CNN - 21 out of 506 slides contained errors of such nature. The other 18 slides were excluded due to markings and other staining errors. After this process, the final dataset contained 467 slides.

### Quantification of spatial features for the segmented tumor tissues

The obtained segmented images from TCGA were then processed further in order to extract spatial information in the form of two types of features: *tissue prevalence* (TIP) and *tumor microenvironment composition* (TMEC). TIP is a distribution of tissue classes within the whole tissue area, i.e. excluding the background class. TMEC measures a distribution of tissues that neighbour the tumor tissue within a predefined margin.

The prevalence *t*_*i*_ of tissue *i* is expressed as:

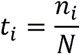

where ni is the number of patches for tissue *i* and *N* is the total number of tissue patches (excluding the background class) and

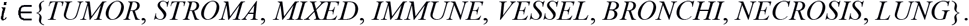

The vector *T* with entries given by ti makes up the TIP features. The background class was omitted, as it’s not relevant to the tissue structure.

The microenvironment composition *m*_*j*_ for tissue *j* is:

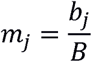

where *b*_*j*_ is the number of patches of class *j* that neighbour the tumor class and *B* is the total number of all patches neighbouring the tumor class (excluding the tumor itself and the background class), with

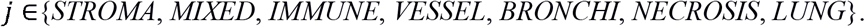

The TMEC features are organized in a vector *M*, with *m*_*j*_ as its entries. The neighbour patches are considered only within a margin around the borders of tumor regions (see **Figure 1b**.). Each tumor patch is considered separately and up to eight neighbours around it are counted. These patches are summed up to *b*_*j*_ for each class *j* and to *B* in total.

**Figure 1.**
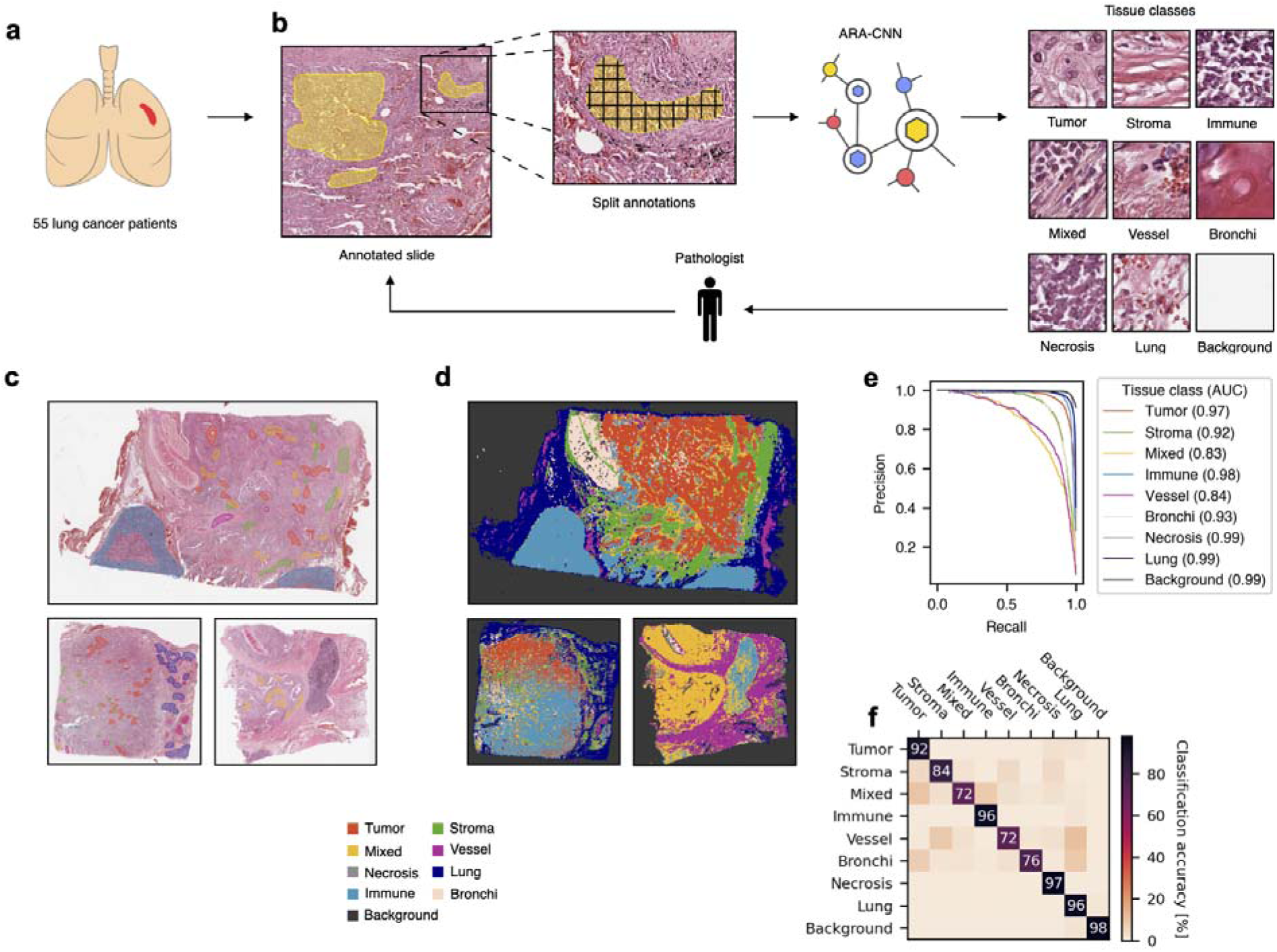
Overview of training ARA-CNN for lung cancer tissue classification **(a)** We sourced H&E tissue slides from 55 lung cancer patients **(b)** These slides were annotated by an expert pathologist in an active learning loop with ARA-CNN, which resulted in the *LubLung* dataset and a trained tissue classification model. **(c)** Example annotations of various tissue regions **(d)** Segmentation results from ARA-CNN show that tissue heterogeneity in the TME is captured correctly **(e)** Precision-recall curves for each tissue class obtained in a 10-fold cross-validation scheme on the *LubLung* dataset. The mean AUC is 0.94. **(f)** Confusion matrix for ARA-CNN trained with *LubLung*. Row labels indicate true classes, while column labels describe classes predicted by the model.

Using the microenvironment and prevalence data, we also calculated three spatial metrics that were previously defined in the literature: intra-tumor lymphocyte ratio (ITLR)^21^, Simpson diversity index^44^, Shannon diversity index^45^. We used a simplified version of these metrics - instead of cell-wise, we calculated them patch-wise. Specifically, these metrics were computed as follows:

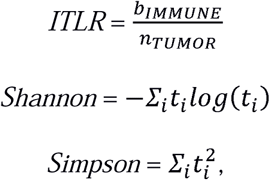

where *b*_*IMMUNE*_ is the number of immune patches that neighbour the tumor and *n*_*TUMOR*_ is the number of tumor patches in the whole slide.

### Mutation classification

The processed data from TCGA served as input in the mutation classification task. The predictor variables consisted of clinical data (age, sex, smoking status, pathologic stage), the previously introduced ITLR, Shannon diversity index, Simpson diversity index, as well as the here proposed TMEC and TIP. The response variables were binary and were defined by the mutation status for the following genes: *ALK, BRAF, DDR2, EGFR, KEAP1, KRAS, MET, PDGFRB, PIK3CA, RET, ROS1, STK11, TP53*.

For each classification task, where the class was specified by the presence of mutation of a given gene, the dataset was oversampled so that positive (mutation occurred) and negative (mutation did not occur) subsets of examples were equal in size. Oversampling was done by inserting multiple copies of the positive examples so that their number reached that of the negative ones.

Several combinations of predictors were tested: clinical, clinical + ITLR, clinical + Shannon diversity index, clinical + Simpson diversity index, clinical + TMEC, clinical + TIP, clinical + TMEC + TIP, TMEC only. To classify the mutation status for each gene, two distinct machine learning models were trained and compared. The first one was a simple linear model in the form of logistic regression. It was fitted using the Liblinear solver^46^, with the L2 penalty and up to 2000 iterations. The second one was the Random Forest algorithm^47^. We used the implementation from the sklearn Python library^48^ with default parameter values.

All models were trained 100 times with 10-fold cross-validation and the resulting classification accuracy metrics were averaged. Classification accuracy was evaluated using the AUC metric.

### Survival prediction data exploration

For the survival prediction task, we defined the set of predictors as the same variables (clinical and spatial) as in the mutation classification task, extended by the set of variables that indicated the mutations in genes. The response variables consisted of the time to the last follow up and the censoring status for the patients. The latter was sourced from TCGA using the curated TCGA Data R package^49^.

We explored the univariate relations between the components of TIP and TMEC and survival, as well as between the gene mutation status and survival, using the Kaplan-Meier estimator. For each tissue type *i*, we compared survival between the groups with high and low *t*_*i*_. Similarly, we compared survival between the groups with high and low *m*_j_. The cutoff point in each case was chosen automatically by the survminer R package. For gene mutations, we divided the patients based on the binary mutation status. The significance of each survival comparison was assessed with the log-rank test.

### Survival modeling

The aforementioned predictors were used as input to the Cox proportional hazards model. They were organized into the following basic variants: clinical, clinical + ITLR, clinical + Shannon diversity index, clinical + Simpson diversity index, clinical + TMEC, clinical + TIP, clinical + TMEC + TIP. In addition, variants with mutation data added on top of clinical data were considered. Each variant was trained in a 10-fold cross-validation schema and for each a resulting median c-index was measured. On top of that, we also plotted hazard ratios for two models with the best c-index. For categorical variables, the hazard ratio of their basal values is set to 1. For the sex variable, the basal value was ‘Female’. For the Stage variable, the basal value was ‘Early stage’. For mutation variables (*EGFR, STK11* and *TP53*) the basal value was the absence of alteration. Finally, for smoking status, non-smoker was set as basal.

## Results

### Implementation and validation of ARA-CNN

To automatically segment H&E images and be able to compute human interpretable spatial features, we first generated a training dataset consisting of whole slide scans of H&E stained tissue sections from 55 LUAD samples from a single institution (**Fig. 1a** and **Materials and methods**). We annotated consistent regions of tissue, marking them as one of the following nine classes: *tumor* with neoplastic epithelial cells; *stroma* composed of connective tissue within tumor or extra-tumoral connective tissue; *mixed* where connective tissue was strongly infiltrated with immune cells; *immune* composed of lymphocytes and plasma cells or fragments of pulmonary lymph nodes; *vessel* composed of smooth muscle layers (veins and arteries) with red blood cells within lumen; *bronchi* composed of cartilage and bronchial mucosa; *necrosis* including necrotic tissue or necrotic debris; *lung* (lung parenchyma); and *background* of the tissue scan (no tissue). We then extracted 87×87 μm tissue patches from each annotated region and used them to iteratively train our previously introduced accurate, reliable and active convolutional neural network (ARA-CNN) model, in each iteration obtaining additional annotations for classes pointed as uncertain by the model^37^ (**Fig. 1b**). We trained the final ARA-CNN on 23,199 patches obtained after three annotation and re-training iterations. We refer to this training dataset as *LubLung* (**Materials and methods**). The TME in the original slides differed between patients, which gave us a diverse set of training examples. A range of tissue classes was selected individually by a pathologist and was used for model training. Some of the slides were more covered by tumor and necrotic cells or stroma, while in others immune infiltration, vessels or mixed class were dominating. In most of the slides we observed the “normal” lung structures, so bronchi was less common and needed more training data from many sections (**Fig. 1c**). The application of the final ARA-CNN on source *LubLung* tissue slides allowed to correctly capture the TME heterogeneity in terms of all trained classes, which was confirmed by a pathologist who compared the original H&E slides with the final output of the model (**Fig. 1d**).

Next, we assessed the classification performance of ARA-CNN. To this end, we used a 10-fold cross-validation procedure on the final set of 23,199 annotated patches obtained in the *LubLung* dataset (**Materials and methods**). The best performance in a single class versus rest classification was achieved for the background, lung, necrosis, tumor, and immune classes (area under the curve, AUC range: 0.97– 0.99) (**Fig. 1e**). The lowest AUC (0.83) was obtained for the mixed class, which is not surprising given that it is a tissue that is a mix of two other classes (stroma and immune). We then computed a confusion matrix, which confirmed that the best trained classes were background, necrosis, lung, immune and tumor (accuracy range: 92.36%–98.01) (**Fig. 1f**). In terms of errors, the model most often confused the mixed class with tumor (9.72% of the patches annotated as mixed were classified as tumor) or immune (8.17% of the patches); the vessel class with stroma or lung (8.73% and 10.79% of the patches, respectively); and the bronchi class with tumor or lung (7.30% and 8.53% of the patches, respectively). Given that patches of these classes were also often hard to distinguish by an expert pathologist, we conclude that our trained ARA-CNN model can reliably classify different tissue types in H&E images of LUAD tissue sections.

### Identification of TME spatial composition features

We then sought to apply our trained ARA-CNN model to study the spatial architecture of the TME in H&E images from 411 LUAD patients downloaded from the TCGA database (**Supplementary Data**). We split each image into 87×87 μm patches and then normalized each patch to the same color space as the images in the *LubLung* dataset (**Materials and methods**). We used each patch as input to our ARA-CNN model, which returned the probabilities of assigning each patch to one of the nine tissue classes. We then segmented each image by assigning the most probable class to each patch. For each image, we computed two metrics reflecting the spatial structure of the TME: TIP and TMEC (**Materials and methods** and **Fig. 2a**). TIP is represented by a vector of values *t*_*i*_, computed as the fraction of patches assigned to class *i* out of all non-background patches in the whole slide image. TMEC is represented by a vector of values *m*_*i*_, computed as the fraction of patches assigned to class *i* out of all non-tumor and non-background tissue types in a predefined margin around the tumor tissue.

**Figure 2.**
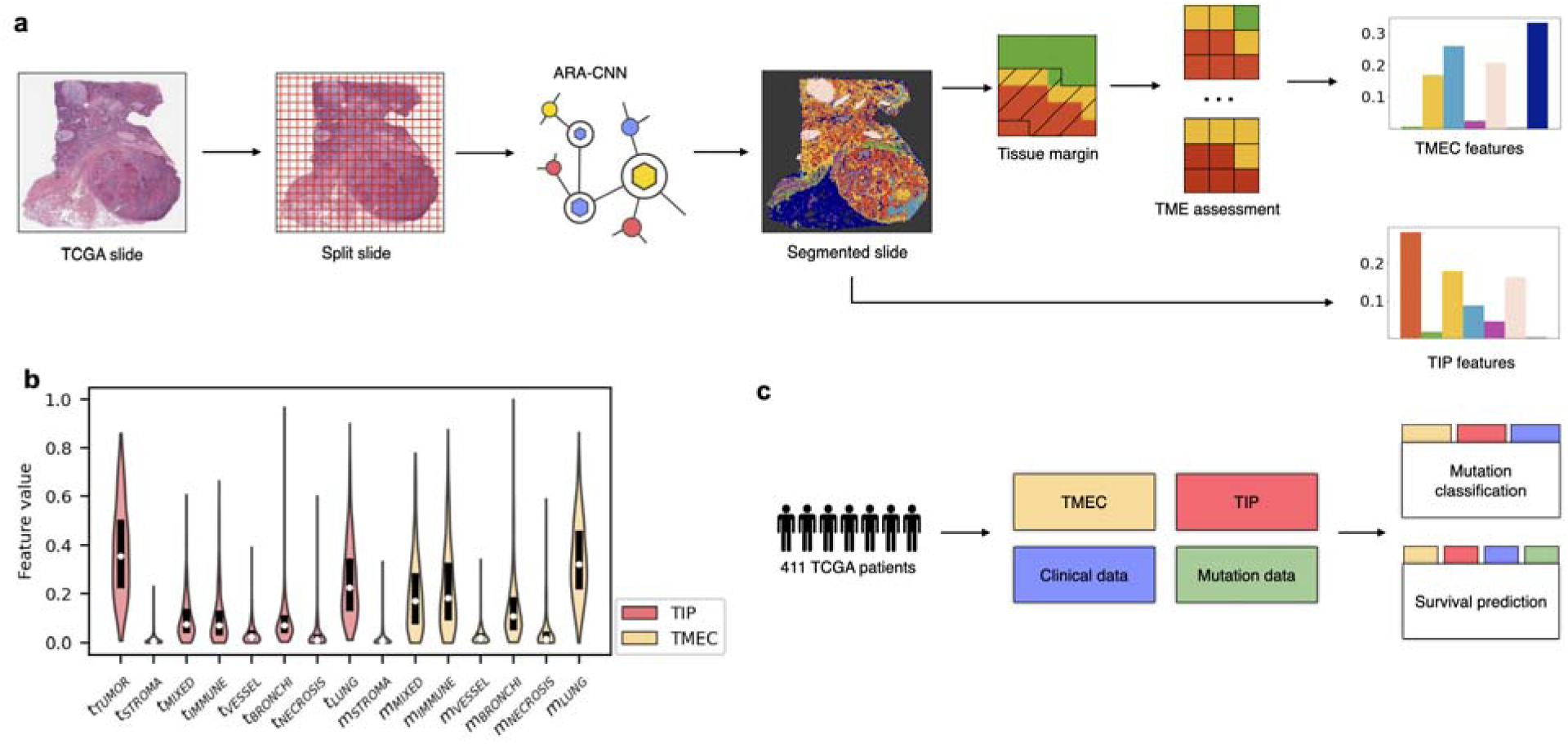
Calculation and utilisation of TIP and TMEC features **(a)** H&E slides from TCGA were downloaded and split into tissue patches. Each patch was classified with ARA-CNN, producing tissue segmentations. These segmentations were next used to calculate the TIP and TMEC features. **(b)** Distribution of individual component features in TIP and TMEC. The most often occurring features for TIP were *t*_*TUMOR*_ and *t*_*LUNG*_. For TMEC, these were *m*_*LUNG*_, *m*_*IMMUNE*_ and *m*_*MIXED*_. **(c)** Tasks performed with the help of the TIP and TMEC features. In addition to the TIP and TMEC features, clinical and mutation data was also sourced from TCGA. These datasets were combined and served as input in two tasks: survival prediction and gene mutation classification. The results were compared to those obtained using previous spatial metrics instead of TIP and TMEC.

Across the investigated tissue classes, tumor and lung classes dominated the entire tissue composition, with a median *t*_*TUMOR*_ of 0.36 and a median *t*_*LUNG*_ of 0.22. The next three most abundant classes in the LUAD slides were mixed, immune and bronchi (with median prevalence of around 0.07). Finally, the least abundant classes were stroma, vessel, and necrosis. The most dominant classes of the tumor microenvironment were lung (median *m*_*LUNG*_ = 0.32), immune (median *m*_*IMMUNE*_ = 0.18) and mixed (median *m*_*MIXED*_ = 0.17). These classes were followed by bronchi (median *m*_*BRONCHI*_ = 0.11). The least abundant in the tumor microenvironment were stroma, vessel and necrosis classes. This indicates that in many patients, the tumor is surrounded by normal lung tissue and is confronted with an immune response. The abundance of all features, however, showed large variability across the analyzed TCGA slides, indicating high heterogeneity of both the entire tissue and the tumor microenvironment composition.

### TME features are predictive of patient survival

We then explored if our metrics can be used to predict patient survival, given that the composition of the TME has been previously shown to influence disease aggressiveness and survival in various cancer types^4,50^. To this end, we first stratified the 411 LUAD patients into two groups based on their TIP and TMEC levels (High vs. Low). For each metric, we compared survival between the two groups using the Kaplan-Meier estimator (**Materials and methods**). Six TIP features (vessel *p* = 0.0016, immune *p* = 0.0058, necrosis *p* = 0.0001, stroma *p* = 0.0352, bronchi *p* = 0.0079 and mixed *p* = 0.0040) and five TMEC features (vessel *p* = 0.0001, immune *p* = 0.0045, necrosis *p* = 0.0009, stroma *p* = 0.0086, and bronchi *p* = 0.0254) showed statistically significant (*p* < 0.05, log rank test, two-sided) differences in survival between High and Low groups (**Fig. 3a-k**). To systematically assess the added value of the TIP or TMEC and to compare them to other predictive features, we trained several versions of a multivariate Cox proportional hazards model of the death hazard for the analyzed LUAD patients and assessed the performance of each model with Harrell’s c-index^51^ (**Materials and methods**). The best model yielded a median c-index of 0.723 and included clinical data (age, sex, pathologic stage, and smoking status), *EGFR, STK11* and *TP53* gene mutations, as well as TIP features, whereas inclusion of TMEC instead of TIP features yielded a slightly lower c-index (0.709) (**Fig. 3l**). All other models – including those based on spatial diversity metrics such as Shannon index^45^, Simpson index^44^ and ITLR (Intra-Tumor Lymphocyte Ratio)^21^ – resulted in lower c-index values. These results indicate that the TIP and TMEC features, which respectively reflect the repertoire of different tissues and their proportions across the entire examined tissue and across the TME,, are superior to other spatial metrics in predicting patient survival.

**Figure 3.**
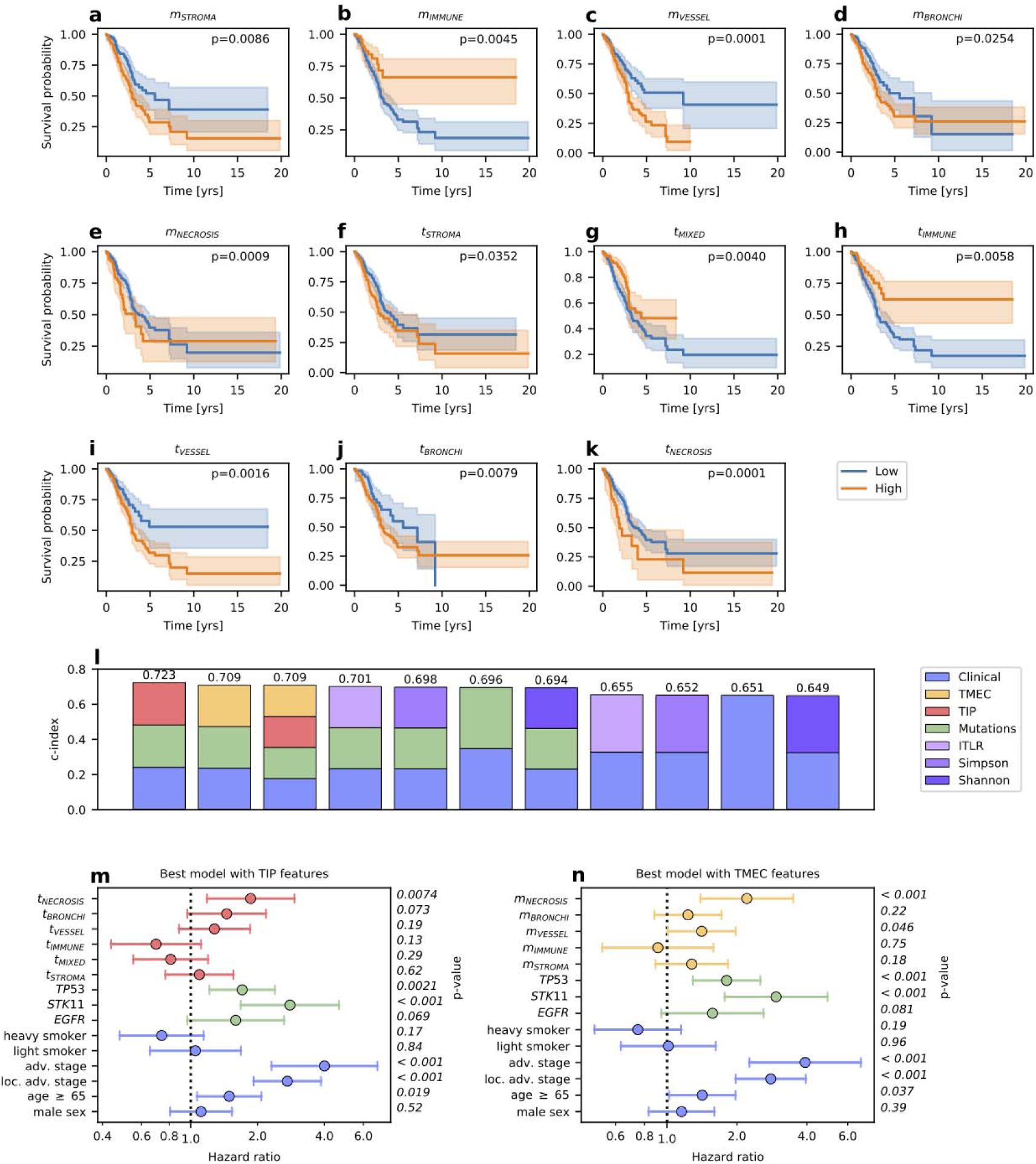
Survival prediction results. **(a-k)** Kaplan-Meier plots for TIP and TMEC features that result in patient stratification into two groups: with high and low values of the feature. Only features with statistically significant differences in patient survival are shown, as measured using the log rank test (p-values in the top right corner). The results correlate with previous studies of the relationship between these features and patient survival. **(l)** c-index scores for Cox models from survival prediction experiments performed with different feature sets. The best results were obtained for models with such feature sets that included TIP and TMEC features. **(m)** Hazard ratios for the best model that utilised the TIP features. The prevalence of the necrosis tissue class in the whole slide has a statistically significant negative effect on survival. **(n)** Hazard ratios for the best model that utilised the TMEC features. The presence of the necrosis tissue class and the vessel tissue class in the TME has a statistically significant negative effect on survival.

Next, we assessed the association between TIP and TMEC features and the death hazard accounting for the context of other features (**Materials and methods**). A hazard ratio of 1 for a given feature indicates that the feature has no effect on survival, whereas a feature with hazard ratio larger than 1 indicates an increased death hazard and, therefore, a negative impact on survival. The best performing model was trained with clinical data, *EGFR, STK11* and *TP53* gene mutations, and TIP features. According to this model, a high abundance of *t_NECROSIS_* and *t_VESSEL_* features in the H&E image was associated with increased hazard. Similarly, abundance of *t_BRONCHI_* and *t_STROMA_* features had a negative effect on survival (**Fig. 3m**). In contrast, *t_IMMUNE_* and *t_MIXED_* features were associated with a decreased death hazard and therefore longer survival (**Fig 3m**), in line with the established role of the immune system as a barrier against tumor progression^4,19,50^. Among mutation features, *TP53* and *STK11* mutations significantly increased (*p* < 0.05, Wald test, two-sided) the death hazard, in agreement with the results of the independent Kaplan-Meier analysis (**Supplementary Fig. 1**). A model trained with TMEC instead of TIP features yielded very similar results (**Fig. 3n**). The impact of clinical features on the hazard agreed with previously published results and our independent Kaplan-Meier analysis (**Supplementary Results**).

### TME features are predictive of disease-relevant mutations

Next, we sought to investigate the association of the human-interpretable spatial composition features of H&E images with mutations in lung cancer genes. To this end, we trained classifiers for the mutation status of 13 genes that are frequently mutated in LUAD: *ALK, BRAF, DDR2, EGFR, KEAP1, KRAS, MET, PDGFRB, PIK3CA, RET, ROS1, STK11, TP53*. We evaluated eight different feature sets (**Materials and methods**) with two machine learning algorithms: logistic regression and random forest. Out of all 104 feature set and gene combinations, logistic regression was the better performing algorithm in 55 cases, while random forest performed better in the remaining 47 cases, indicating that for some genes non-linear relationships between the predictive features may be relevant for prediction of their mutations (**Table 1**). For 8 out of 13 considered genes (namely, *RET, KRAS, KEAP1, TP53, BRAF, PDGFRB, ROS1, STK11*), using the TIP or TMEC features gave the best result. For the remaining 5 genes (*MET, ALK, DDR2, PIK3CA, EGFR*), the best AUC was reached for models that utilized one of previously existing spatial metrics as features.

**Table 1.** Mutation/rearrangement classification AUC scores (given as % of area under the precision-recall curve) for TCGA LUAD patients. The best result for each gene is marked in bold. In cases where the random forest classifier gave the best result, the cells are colored in yellow. Otherwise, if logistic regression gave the best result, the cells are colored in light blue.

| Gene | Mutation count | Clinical | Clinical + ITLR | Clinical + Shannon | Clinical + Simpson | Clinical + TMEC | Clinical + TIP | Clinical + TMEC + TIP | TMEC |
| --- | --- | --- | --- | --- | --- | --- | --- | --- | --- |
| *RET* | 25 | 53.70 ± 2.21 | 50.76 ± 2.39 | 58.03 ± 2.13 | 57.76 ± 2.09 | **67.14 ± 2.75** | 59.46 ± 2.13 | 65.32 ± 1.95 | 64.39 ± 2.73 |
| *KRAS* | 96 | 61.12 ± 0.88 | 61.36 ± 0.84 | 60.06 ± 1.04 | 60.51 ± 0.99 | **62.37 ± 1.50** | 62.07 ± 1.66 | 60.98 ± 1.71 | 55.58 ± 1.54 |
| *KEAP1* | 96 | 51.35 ± 1.12 | 53.98 ± 1.60 | 51.37 ± 1.17 | 50.87 ± 1.21 | **61.07 ± 1.60** | 60.33 ± 1.85 | 60.05 ± 1.58 | 56.85 ± 1.48 |
| *TP53* | 200 | 55.28 ± 0.74 | 54.38 ± 0.75 | 55.18 ± 0.78 | 55.02 ± 0.79 | **58.59 ± 1.30** | 57.87 ± 1.26 | 57.78 ± 1.52 | 57.14 ± 1.48 |
| *BRAF* | 30 | 55.38 ± 1.74 | 55.14 ± 2.49 | 56.59 ± 1.76 | 56.87 ± 1.74 | 55.26 ± 2.66 | **57.10 ± 1.95** | 56.23 ± 2.14 | 51.35 ± 2.88 |
| *PDGFRB* | 15 | 67.85 ± 2.51 | 69.75 ± 2.79 | 69.91 ± 3.23 | 70.06 ± 2.97 | 72.03 ± 2.04 | 72.78 ± 2.48 | **73.50 ± 2.02** | 52.13 ± 3.83 |
| *ROS1* | 18 | 58.29 ± 3.58 | 60.74 ± 2.76 | 51.27 ± 3.20 | 49.97 ± 3.20 | 59.44 ± 4.69 | 60.07 ± 2.55 | **61.45 ± 4.54** | 59.58 ± 4.08 |
| *STK11* | 44 | 53.98 ± 1.76 | 53.91 ± 1.67 | 53.56 ± 1.83 | 61.39 ± 2.22 | 58.41 ± 1.51 | 54.89 ± 2.64 | 59.22 ± 2.30 | **62.89 ± 2.51** |
| *MET* | 19 | 61.22 ± 2.31 | **69.20 ± 2.31** | 63.14 ± 2.39 | 64.19 ± 2.35 | 66.58 ± 3.70 | 68.10 ± 2.45 | 63.68 ± 2.76 | 63.36 ± 4.28 |
| *ALK* | 28 | 53.55 ± 2.23 | 48.42 ± 2.49 | **62.64 ± 3.05** | 59.51 ± 2.23 | 50.32 ± 2.88 | 61.53 ± 1.70 | 58.71 ± 1.95 | 49.41 ± 2.89 |
| *DDR2* | 18 | 57.17 ± 3.81 | 56.89 ± 4.21 | **57.38 ± 3.80** | 56.69 ± 3.82 | 55.63 ± 4.30 | 54.65 ± 3.96 | 52.56 ± 4.37 | 51.78 ± 5.13 |
| *PIK3CA* | 23 | 49.14 ± 1.78 | 51.30 ± 2.67 | 51.07 ± 2.11 | **58.35 ± 2.78** | 53.90 ± 3.27 | 56.96 ± 3.46 | 58.01 ± 3.18 | 56.15 ± 3.22 |
| *EGFR* | 49 | **65.09 ± 1.50** | 61.48 ± 2.49 | 63.32 ± 1.25 | 64.90 ± 1.97 | 59.01 ± 1.60 | 59.69 ± 1.56 | 57.93 ± 1.60 | 49.05 ± 1.97 |

The best AUC (73.5%) was reached for the *PDGFRB* gene mutation by a classifier using clinical data and both TIP and TMEC as features (**Table 1**). The best model without TIP and TMEC, and with the Simpson metric as a feature, yielded an AUC smaller by 3.4 percentage points (p.p.). This shows that for the *PDGFRB* gene mutation, the full information about tissue distribution, not reduced to a single value using entropy and without focusing on only selected tissues, is highly relevant for its mutation status. The classification performance of the best model using both TIP and TMEC for that gene is only slightly smaller than AUC of 75%, as previously reported for a deep learning model trained on raw H&E images^34^, but is less difficult to interpret. For eight other genes (*RET, KRAS, KEAP1, ROS1, STK11, MET, ALK, EGFR*), the best AUC ranged between 60% and 70%, while for the four remaining ones (*TP53, BRAF, DDR2, PIK3CA*) the best AUC ranged between 55% and 60%. For some of the genes, the inclusion of TIP or TMEC features resulted in impressive improvements compared to other feature sets. For *RET*, the model trained with clinical data and TMEC outperformed the best model without TIP and TMEC features, but including the Shannon metric, by around 9.1 p.p. Similarly, for *KEAP1* the classification performance increased by 7 p.p. compared to models without TIP or TMEC. These results indicate that, in LUAD, there exists a subset of tumor mutations that significantly correlate with how the TME is structured, and that both TIP and TMEC features are predictive of the presence of these mutations.

We then inspected the two best performing models in the mutation classification task that utilised TIP and TMEC features to find which predictor features were the most important for identifying mutations. Both of the algorithms used – logistic regression and random forest – are easily interpretable because they allow effective identification of the most important features. First, we analysed the logistic regression classifier of *PDGFRB* mutations with clinical, TMEC and TIP features (**Fig. 4a**). The most important features positively correlated with *PDGFRB* mutation were sex, *m*_*MIXED*_ – corresponding to the proportion of the mixed tissue in the tumor microenvironment – and *t*_*TUMOR*_ – corresponding to the fraction of the entire slide occupied by the tumor. On the other hand, the most negatively correlated (i.e., decreasing the chance of mutation) features were non-smoker status, *t*_*IMMUNE*_, and *m*_*BRONCHI*_. Next, we inspected the random forest classifier of *RET* mutations, which included clinical and TMEC features in its feature set (**Fig. 4b**). The latter proved to be of larger importance than the former ones. Indeed, *RET* mutations were found to be most associated with the prevalence of different tissues in the tumor microenvironment, with bronchi and vessels identified as the most impactful tissues, followed by mixed, stroma, lung, immune and necrosis. This observation might be explained by the fact that, in LUAD, *RET* mutations mainly consist of rearrangements between *RET* gene and its common fusion partners such as *KIF5B, CCDC6, CUX1, TRIM33, NCOA4, KIAA1468* and *KIAA1217* genes.

**Figure 4.**
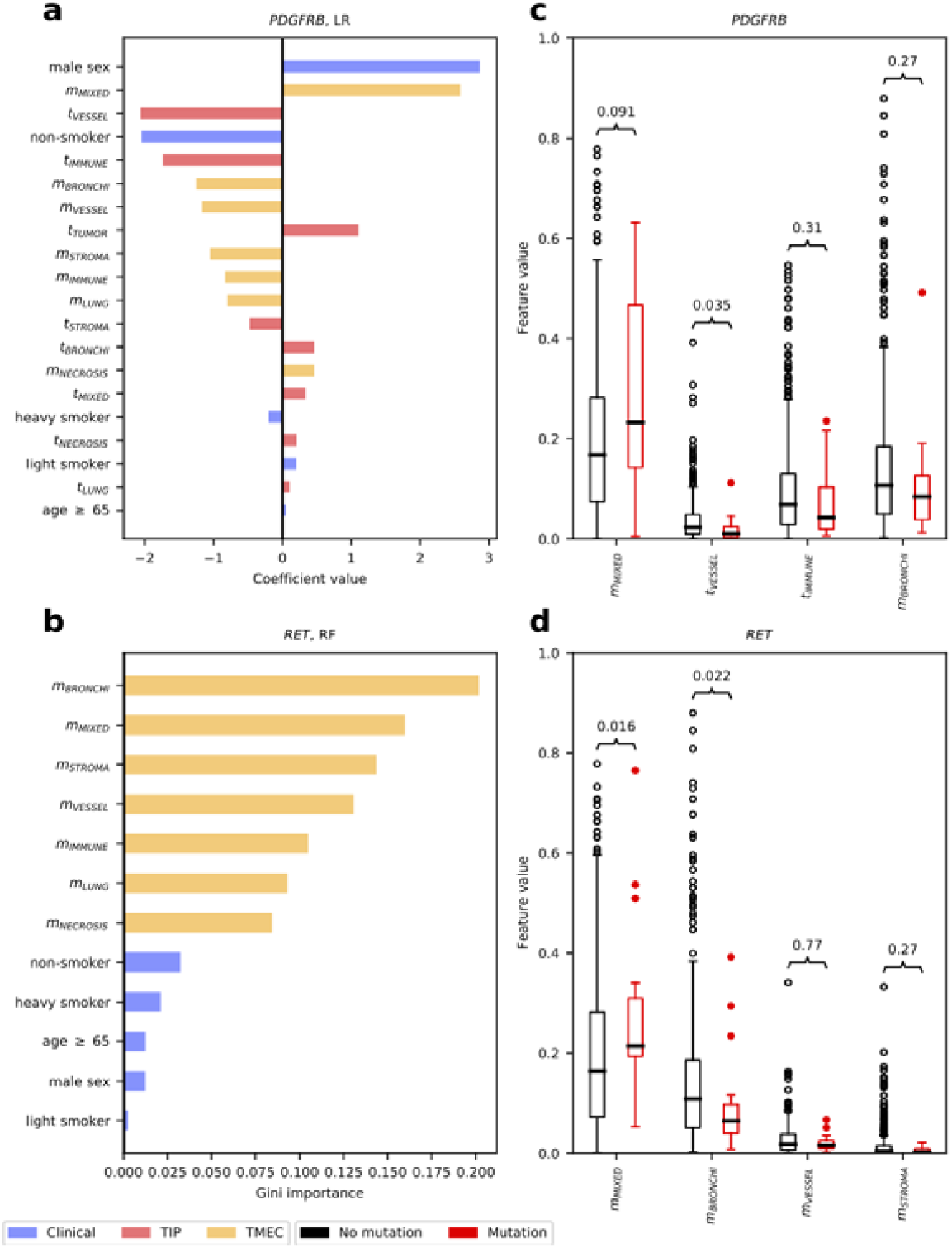
Feature importance for the two best performing mutation classification models that utilised TIP and TMEC features **(a)** Feature importance for the *PDGFRB* gene mutation classifier (logistic regression). Here, feature importance is measured by the value of its regression coefficient. **(b)** Feature importance for the *RET* gene mutation classifier (random forest). Here, the importance is measured by the reduction of the Gini index obtained when the feature is added to the tree, averaged across the trees in the random forest model. **(c)** Distribution of feature values for four of the most important TIP or TMEC features, as presented in (a), divided between patients with the mutated and non-mutated *PDGFRB* gene. **(d)** Distribution of feature values for four of the most important TMEC features, as presented in (b), divided between patients with the mutated and non-mutated *RET* gene.

In addition to feature importance, we also inspected the distributions of the values of the TIP and TMEC spatial composition features for patients with and without mutations of the *PDGFRB* and *RET* genes. For both of them, we selected the four most important TIP or TMEC features and assessed their value distributions separately for mutated and non-mutated cases. For *PDGFRB*, these features were: *m*_*MIXED*_ and *m*_*BRONCHI*_ (TMEC features), as well as *t*_*VESSEL*_ and *t*_*IMMUNE*_ (TIP features) (**Fig. 4c**). We detected a statistically significant difference between the value distributions (two-sided Wilcoxon test *p-value* < 0.05) for *t*_*VESSEL*_. For *RET*, the four most important features were TMEC features *m*_*BRONCHI*_, *m*_*MIXED*_, *m*_*VESSEL*_ and *m*_*STROMA*_ with *m*_*MIXED*_ and *m*_*BRONCHI*_ features having a statistically significant difference in value distributions between mutated and non-mutated tumors (**Fig. 4d**). These results indicate that the spatial composition features TIP and TMEC are different between tumors with and without *PDGFRB* and *RET* mutations, and their importance for the classification of mutations of these genes is not incidental.

## Discussion

We have developed a novel H&E image classification model, ARA-CNN, and a training dataset of annotated tissue patches from LUAD H&E images, *LubLung*. Both considerably expand the current ability to analyze the TME automatically and quantitatively in lung cancer samples, which in turn has important implications for patient stratification and precision treatment. TIP and TMEC metrics, which we have introduced in this work, provide a novel way of capturing the composition and spatial structure of the TME, and are predictive of both overall survival and clinically relevant mutations. Spatial statistics of H&E images in the form of metrics that quantify colocalization of cell or tissue types, have been previously shown to be predictive of patient survival^21^. However, these metrics are computed based on a limited number of features, such as counts of tumor and immune cells. Other approaches that try to link the structure of tumor tissue and TME with either gene mutations or patient survival are end-to-end deep learning models and work as ‘black boxes’^29,30,34,52–54^. Instead, our approach allows explicit interpretability, as it decouples H&E slide inference from downstream tasks (*e*.*g*., mutation classification and survival analysis). The TIP and TMEC features are *per se* human interpretable, so it is possible to precisely pinpoint which tissue types are the most important. Our approach requires the initial tissue classification to be as accurate as possible. We ensured this to be the case by using ARA-CNN, which performs excellently in classifying nine tissue classes present in lung cancer H&E images. To foster further research in predictive spatial statistics based on a rich repertoire of segmented lung cancer tissues, in addition to *LubLung* we also share the segmented TCGA images as a separate dataset, named *SegLungTCGA*.

Our analysis revealed that patient stratification based on TIP and TMEC features yields significant differences in patient survival between the strata. Moreover, the most predictive survival models included TIP and TMEC features. These findings are supported by previous clinical studies. It has been shown that blood vessel invasion is a major prognostic factor in lung cancer survival^14,55^. Similarly, there have been studies which proved that tumor necrosis is a significant risk factor for survival in lung cancer^56^. However, the complexity of the entire lung microenvironment plays a key role in the development of primary lung carcinomas and offers a resource of targets for personalized therapy development. Targeting the angiogenesis and immune cells has elucidated the prognostic and pathophysiological roles of other components of the TME in lung cancer^13,57^. In the end, the combination of the clinical and genetic information with the TME landscape may play a pivotal role in predicting the type and duration of response to personalized therapies.

We found eight genes relevant to lung cancer (*PDGFRB, RET, KRAS, KEAP1, ROS1, STK11, MET* and *ALK*), for which integrating clinical data with our TME metrics clearly improves the ability to predict mutations in these genes. We speculate that mutations of these genes may alter cellular interactions, and hence the spatial arrangement of the TME visible in H&E images. For *RET, ROS1* and *ALK* genes, mutations mainly consist of chromosomal rearrangements which produce chimeric proteins that might affect the cellular organisation within the TME^58–60^. Likewise, loss of *STK11/LKB1* overlapping with oncogenic *KRAS* mutations is associated with increased neutrophil recruitment, and decreased T-cells infiltration in lung cancer tumors^61^. Moreover, *STK11* mutations often coexist with *KEAP1* mutations that relate to cellular resistance to oxidative stress^62^, and co-occurrence of *KEAP1* mutations and *PTEN* inactivation is an indicator of an immunologically “cold” tumor^63^. We speculate that each of these mutations might slightly affect the cellular morphology in H&E images in a way that is not apparent to the human eye, but can be captured by deep-learning algorithms.

Our findings concern mutations of clinically relevant genes, and as such may have clinical implications. For example, both *RET* and *PDGFRB* are clinically relevant LUAD cancer genes. *RET* has proto-oncogene properties and its fusions, which occur in 1–2% of LUAD^64^, are associated with a high risk of brain metastasis^65^. However, last clinical trials indicated that they may be effectively targeted by RET tyrosine kinase inhibitors as pralsetinib, selpercatinib^64^. *PDGFRB* is a member of the PDGF/PDGFR axis that is recognized as a key regulator of mesenchymal cell activity in TME^66^, and several new agents (linifanib, motesanib, olaratumab) that block the PDGFR signaling are being tested in LUAD^67^. In breast, colon, pancreas and prostate cancers, the high stromal expression of the PDGFRβ protein has been associated with poor prognosis^67^, however its prognostic relevance in tumors of epithelial origin is inconclusive^66^. It was only confirmed that a relative expression of PDGFRs is a strong and independent predictor of longer survival for surgical stages of lung cancer (I-IIIA)^67^.

Our approach has several limitations. In the mutation classification tasks, we used simple machine learning models – logistic regression and random forest. An end-to-end deep learning model might give better results, however, as discussed above, these models suffer when it comes to interpretability. Another limitation is the fact that ARA-CNN works on a patch-based basis. An alternative to that is a cell-based classifier, which could produce more fine-grained segmentations and in turn enable a more precise computation of spatial statistics. On the other hand, with a patch-based approach, a suitably small patch size can be selected, as we did in this study. Such small patches can be assumed to be homogeneous when it comes to cell types and can enable a precise computation of summary statistics such as the TIP and TMEC metrics, that we have introduced here. Furthermore, in the survival prediction task we did not have access to and hence did not utilise the treatment information as features. Since treatment has a large impact on survival, using this data is expected to improve prediction performance by a large margin. However, the main focus of our study was not to deliver the best performing survival prediction approach, but rather to assess the predictive power of our proposed spatial composition features, which can be performed without including the treatment data in the model. Lastly, to apply our approach to another cancer type, one would need to retrain the ARA-CNN model, which necessitates substantial input from a trained pathologist. This training effort can be minimised by utilising the active learning component of ARA, which shortens the number of iterations required to build an effective training dataset. For colorectal cancer, a pre-trained model is available from a previous study^37^.

The analysis presented here shows that there is a correspondence between the spatial structure in H&E images for LUAD and both gene mutations and patient survival. Not every mutation is expected to have an effect on tissue prevalence or tumor neighbourhood structure, so it is not surprising that for some of the analyzed genes the mutation classification performance did not exceed an AUC of 0.6. In contrast, it is striking that there are genes for which adding tissue composition data to the clinical information improves classification results. Finally, it is also surprising that our TIP and TMEC metrics, as well as other metrics of TME spatial organization, such as ITLR, can give good results in terms of both mutation classification and survival analysis.

We have explored the applicability of human interpretable features extracted from deep learning-based segmentation of H&E images. Our approach can provide important insights for designing novel cancer treatments, by linking the spatial structure of the tumor microenvironment in LUAD to gene mutations and patient survival. It can also expand our understanding of the effects that the tumor microenvironment has on tumor evolutionary processes. We therefore envision that, in the future, our quantitative approach will become incorporated in routine diagnostics for LUAD and other cancer types.

## Data availability

The *LubLung* dataset is available publically at github.com/animgoeth/LubLung

The *SegLungTCGA* dataset is available at github.com/animgoeth/SegLungTCGA

The ARA-CNN weights trained on *LubLung* are available at github.com/animgoeth/ARA-CNN, in the ‘pretrained’ directory.

The Supplementary Data is available per authors’ discretion.

## Author contributions

Ł.R. performed all experiments, analysed the results and prepared visualisations. I.P. was responsible for H&E image annotation and segmentation validation. Ł.K., I.P., M.N. and E.S. coordinated the active learning process. M.K. contributed to the work on survival analysis. M.B. provided code for extracting mutation data from TCGA. M.N. and P.K. helped with the interpretation of clinical data and provided feedback on the results. J.S. provided the access for the archival tissue material used for the model training. M.N collected the material and prepared the tissue sections. T.K performed the H&E staining and scanned all the slides used for the model training. E.S. and N.C. supervised the research. Ł.R. and E.S. wrote the manuscript. Ł.R., E.S., M.N. and N.C. conceptualised the project. All authors reviewed the manuscript.

## Corresponding author

Correspondence should be addressed to Ewa Szczurek.

