## Supplementary Information for "Deep learning-based tumor microenvironment segmentation is predictive of tumor mutations and patient survival in non-small-cell lung cancer"

#### Abstract

This supplementary information includes **Supplementary Methods**, **Supplementary Results**, **Supplementary Figure 1** and **Supplementary Tables 1-3**.

#### Supplementary Results

**Clinical features identified as significant for patient survival**

From among clinical features, statistically significant association was found for the advanced stage, locally advanced stage and age ≥ 65 features (Wald test *p-value* < 0.05). As expected, both for patients with locally advanced stage and with advanced stage the hazard was increased compared to early stage, and with the advanced stage having the largest estimated hazard ratio of the stages. Similarly, as expected, for older patients  the hazard was larger. The estimated hazard ratio for light smokers was increased as compared to the non-smokers. Seemingly counter-intuitively, the hazard ratio for heavy smokers decreased. This may be explained by the fact that smoking is a well known risk factor for morbidity in lung cancer, but not mortality. In fact, patients who smoke after lung cancer diagnosis have a worse prognosis than patients who quit smoking. However, patients who smoke heavily before diagnosis may  have a high tumor mutation burden, a high number of tumor antigens, and an active immune system with higher sensitivity to immune checkpoint inhibitors, which may prove beneficial for overall survival[1]. From among mutation features, *TP53* and *STK11* statistically significantly increased the death hazard (Wald test *p-value* < 0.05), which agrees with the results from the independent Kaplan-Meier analysis (**Sup. Fig. 1**).

### Supplementary Methods

**Hematoxylin and eosin staining**

FFPE tissue was cut into 3 μm fragments and placed on glass slides. The slides were then stained with hematoxylin and eosin (H&E) in Leica Autostainer XL device (Leica Biosystems, USA), with pre-staining xylene and ethanol wash steps and 7 min of hematoxylin, and 20 sec of eosin. Post-staining wash steps were also included in the program. The glass slides were cover-slipped directly after the staining procedure. The H&E stained slides were scanned using the Aperio ScanScope CS2 device (Leica Biosystems, USA) equipped with an 20x Olympus microscope lens. The images were stored on an internal server in the form of SVS files, which were further analyzed.

**TCGA data extraction and processing**

Independent patient data, including H&E images, mutations, and clinical information was extracted from the TCGA database (up to date as of 2020-05-14). The database contained 478 lung adenocarcinoma (LUAD) cancer patients with H&E tissue slide data. Out of these, frozen tissue slides were filtered out, which left us with 514 images. These slides were downloaded through a REST API provided by TCGA. We developed a parallelized pipeline that downloaded the slides, ran all necessary calculations and then removed the processed images.

Each processed slide was split into non-overlapping patches with side size of 87 μm, same as for the patches in *LubLung*. As an optimisation step, we filtered the extracted patches and excluded the ones where most of the area was empty. To perform this filtering, we first converted each patch to grayscale using the standard Rec. 601 luma formula. Then, we mapped each pixel to either black (pixel value < 200) or white (pixel value ≥ 200) and counted them. The patch was deemed as relevant if the proportion of black pixels to white pixels was larger than 0.05.

In addition to the slides, clinical and mutation data was also extracted from TCGA. The former was downloaded with the curatedTCGAData R package[2] and it contained data for 518 LUAD patients. However, we removed data for Asian patients, as they are noted to be very distinct when it comes to disease progression[3,4]. This left us with clinical data for 510 patients. Mutation data for 563 LUAD patients was downloaded from the UCSC Xena Browser[5] (dated 07-20-2019), in the form of a *TCGA-LUAD.muse_snv.tsv* file. In addition, it was downsized to include only genes selected as relevant to lung cancer.

All datasets were merged together by the TCGA patient identifier. This left us with an intersection between the image, clinical and mutation datasets, which contained 444 patients and 506 slides.

As a last step, the clinical variables were pre-processed in the following manner. Age was quantified into two groups: 65 years and older, younger than 65 years. Sex was set to 1 for male and 0 for female patients. Pack years were quantified into three groups: non-smoker (0 pack years or smoking history set as ‘lifelong non-smoker’ or if pack years were missing, smoking history set to ‘current reformed smoker for > 15 years’), light smoker (less than 30 pack years or if pack years were missing, smoking history set as ‘current reformed smoker for < or = 15 years’) and heavy smoker (30 or more pack years and smoking history set to ‘current smoker’). Pathologic stage was mapped into three groups as well: early (stage I, Ia, Ib), locally advanced (stage II, IIa, IIb, IIIa) and advanced (stage IIIb, IV).

Due to the fact that the tissue slides stored in TCGA exhibit high color variation, they needed to be normalized to a common color space, matching that of the training dataset. Three normalization algorithms were considered: Reinhard et al.[6], Macenko et al.[7] and Vahadane et al.[8]. To decide which of these three should be used on the TCGA data, a series of experiments was conducted, in which the *LubLung* training dataset was normalized with each of these algorithms and then ARA-CNN was trained in a cross-validation schema. The results showed that the best classification performance (mean accuracy 81.52% for Macenko et al., 82.39% for Vahadane et al., 85.76% for Reinhard et al.) was achieved for the dataset variant normalized with the Reinhardt et al. algorithm.

Each relevant patch extracted from the slides was normalized individually with the Reinhardt et al. procedure. The normalization was performed with a region of interest image selected at random from the training dataset. All image patches from the TCGA database were transformed to match the color space of that image.

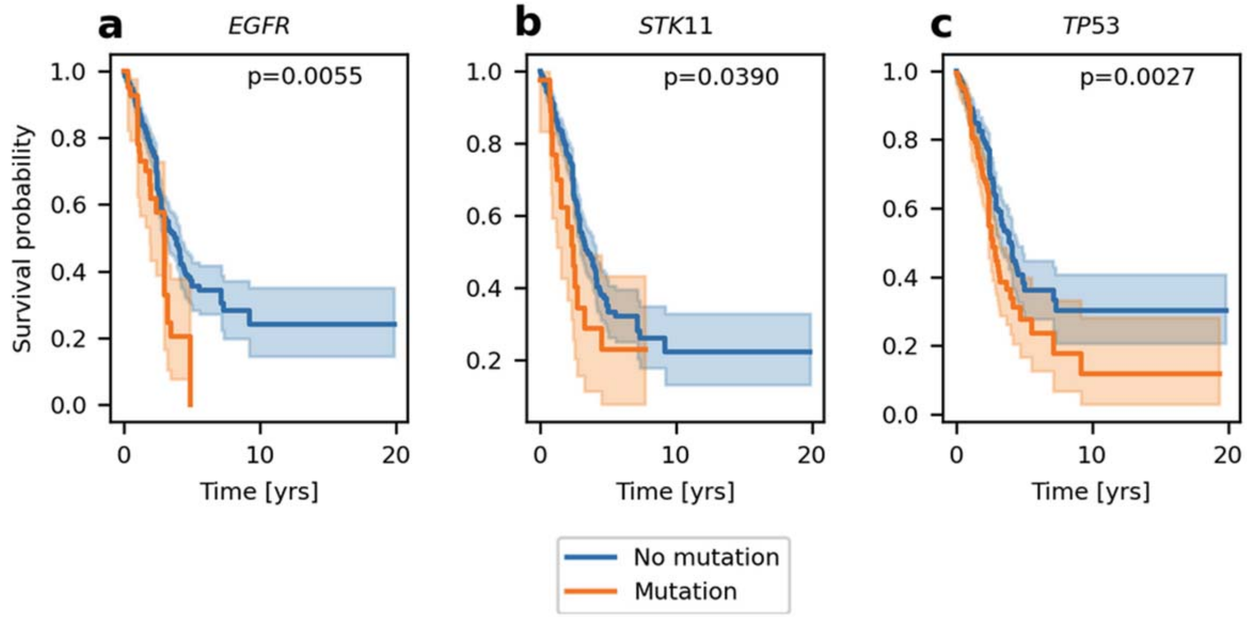


**Supplementary Figure 1.** Kaplan-Meier plots for EGFR, STK11 and TP53 genes, stratified into patients with mutation and patients without mutation. The p-values were measured using the log rank test.

**Supplementary Table 1.** Confusion matrix for ARA-CNN trained with patches sized 74 µm extracted from initial tissue annotations. The mean accuracy is 84.35%.

| | Tumor [%] | Stroma [%] | Mixed [%] | Immune [%] | Vessel [%] | Bronchi [%] | Necrosis [%] | Lung [%] | Background [%] |
|---|---|---|---|---|---|---|---|---|---|
| **Tumor** | 90.34 | 0.25 | 1.15 | 3.72 | 0.59 | 0.86 | 0.88 | 2.21 | 0.00 |
| **Stroma** | 4.17 | 85.37 | 2.66 | 0.19 | 2.36 | 0.77 | 3.40 | 1.08 | 0.00 |
| **Mixed** | 14.65 | 6.73 | 61.49 | 10.69 | 1.09 | 0.99 | 1.68 | 2.67 | 0.00 |
| **Immune** | 0.59 | 0.32 | 0.48 | 94.89 | 0.59 | 0.16 | 0.65 | 2.15 | 0.16 |
| **Vessel** | 1.21 | 11.66 | 1.41 | 0.25 | 70.70 | 2.36 | 1.51 | 10.80 | 0.10 |
| **Bronchi** | 9.93 | 2.37 | 2.63 | 2.09 | 2.73 | 68.85 | 0.83 | 10.58 | 0.00 |
| **Necrosis** | 0.58 | 0.80 | 0.42 | 0.83 | 1.25 | 0.11 | 95.50 | 0.51 | 0.00 |
| **Lung** | 0.32 | 0.10 | 0.04 | 0.01 | 0.38 | 0.20 | 0.02 | 98.31 | 0.61 |
| **Background** | 0.00 | 0.00 | 0.00 | 0.00 | 0.12 | 0.00 | 0.00 | 3.59 | 96.29 |

**Supplementary Table 2.** Confusion matrix for ARA-CNN trained with patches sized 87 µm extracted from initial tissue annotations. The mean accuracy is 85.21%.

| | Tumor [%] | Stroma [%] | Mixed [%] | Immune [%] | Vessel [%] | Bronchi [%] | Necrosis [%] | Lung [%] | Background [%] |
|---|---|---|---|---|---|---|---|---|---|
| **Tumor** | 90.27 | 0.27 | 0.91 | 4.17 | 0.73 | 0.65 | 1.08 | 1.94 | 0.00 |
| **Stroma** | 4.56 | 84.31 | 2.06 | 0.06 | 4.69 | 0.56 | 3.31 | 0.44 | 0.00 |
| **Mixed** | 15.32 | 4.19 | 62.58 | 10.32 | 3.23 | 0.97 | 1.61 | 1.77 | 0.00 |
| **Immune** | 0.58 | 0.08 | 0.58 | 94.79 | 0.74 | 0.08 | 0.58 | 2.15 | 0.41 |
| **Vessel** | 0.80 | 9.73 | 1.47 | 0.20 | 75.07 | 1.60 | 1.67 | 9.40 | 0.07 |
| **Bronchi** | 8.32 | 1.42 | 2.84 | 2.53 | 2.68 | 69.21 | 0.63 | 12.32 | 0.05 |
| **Necrosis** | 0.79 | 0.52 | 0.23 | 0.47 | 0.77 | 0.09 | 96.82 | 0.32 | 0.00 |
| **Lung** | 0.53 | 0.88 | 0.08 | 0.13 | 1.19 | 0.45 | 0.20 | 95.78 | 0.76 |
| **Background** | 0.00 | 0.00 | 0.00 | 0.00 | 0.00 | 0.00 | 0.00 | 1.95 | 98.05 |

**Supplementary Table 3.** Confusion matrix for ARA-CNN trained with patches sized 100 µm extracted from initial tissue annotations. The mean accuracy is 84.35%.

| | Tumor [%] | Stroma [%] | Mixed [%] | Immune [%] | Vessel [%] | Bronchi [%] | Necrosis [%] | Lung [%] | Background [%] |
|---|---|---|---|---|---|---|---|---|---|
| **Tumor** | 88.22 | 0.44 | 1.63 | 3.89 | 0.59 | 1.33 | 1.56 | 2.33 | 0.00 |
| **Stroma** | 6.58 | 80.17 | 2.05 | 0.34 | 3.68 | 1.37 | 4.87 | 0.94 | 0.00 |
| **Mixed** | 12.83 | 4.78 | 60.65 | 11.52 | 1.74 | 1.74 | 3.48 | 3.26 | 0.00 |
| **Immune** | 0.88 | 0.25 | 0.63 | 95.38 | 1.00 | 0.00 | 0.50 | 1.38 | 0.00 |
| **Vessel** | 1.43 | 9.81 | 0.76 | 0.29 | 74.19 | 1.62 | 2.57 | 9.14 | 0.19 |
| **Bronchi** | 9.04 | 1.76 | 1.76 | 2.35 | 2.65 | 67.87 | 0.96 | 13.60 | 0.00 |
| **Necrosis** | 0.60 | 0.40 | 0.23 | 0.63 | 0.80 | 0.13 | 97.00 | 0.20 | 0.00 |
| **Lung** | 0.36 | 0.59 | 0.02 | 0.13 | 0.43 | 0.27 | 0.34 | 97.15 | 0.70 |
| **Background** | 0.00 | 0.00 | 0.00 | 0.00 | 0.00 | 0.00 | 0.00 | 1.47 | 98.53 |
